# Multi-organ transcriptomic landscape of *Ambystoma velasci* metamorphosis

**DOI:** 10.1101/2020.02.06.937896

**Authors:** Palacios-Martínez Janet, Caballero-Pérez Juan, Espinal-Centeno Annie, Marquez-Chavoya Gilberto, Lomelí Hilda, Salas-Vidal Enrique, Schnabel Denhi, Chimal-Monroy Jesus, Cruz-Ramírez Alfredo

## Abstract

Metamorphosis is a postembryonic developmental process that involves morphophysiological and behavioral changes, allowing organisms to adapt into a novel environment. In some amphibians, aquatic organisms undergo metamorphosis to adapt in a terrestrial environment. These organisms experience major changes in their circulatory, respiratory, digestive, excretory and reproductive systems. We performed a transcriptional global analysis of heart, lung and gills during diverse stages of *Ambystoma velasci* metamorphosis. In our analyses, we identified eight gene clusters for each organ, according to the expression patterns of differentially expressed genes. We found 4,064 differentially expressed genes in the heart, 4,107 in the lung and 8,265 in the gills. Among the differentially expressed genes in the heart, we observed genes involved in the differentiation of cardiomyocytes in the interatrial zone, vasculogenesis and in the maturation of coronary vessels. In the lung, we found genes differentially expressed related to angiogenesis, alveolarization and synthesis of the surfactant protein. In the case of the gills, the most prominent biological processes identified are degradation of extracellular matrix, apoptosis and keratin production. Our study sheds light on the transcriptional responses and the pathways involved in the transformation of the facultative metamorphic salamander *A. velasci* in an organ-specific manner.

## Introduction

Metamorphosis is a dynamic postembryonic developmental process that involves changes in morphophysiology of the transformed organism. This process is widely distributed in many invertebrate and vertebrate groups^1^ such as cnidarians, insects, fishes and amphibians. During metamorphosis, a series of changes take place to allow the organism to adapt in different environments^2,3,4,5^. In amphibians, this process allows aquatic tadpoles to transform into terrestrial juvenile organisms^1,6,7,16^. This metamorphosis process is under the control of thyroid hormones (THs)^1^. Once liganded with THs, the thyroid hormone receptor-retinoic acid receptor (TR-RXR) heterodimers recruit coactivators and induce the transcription of its target genes, which in turn direct the cellular metamorphic reprogramming events^1,6,7^.

Metamorphic functional changes occur in the nervous, respiratory, circulatory, digestive, integumentary and reproductive systems^6,8^. The changes involve pivotal cellular and tissular processes such as cell differentiation, cell proliferation, programmed cell death and tissue remodeling^4,9,10,15^. For example, in *Xenopus laevis* metamorphosis, in the tail, gills and interdigital tissue the process that predominates is tissue remodeling^11^, while in the nervous and digestive systems the predominant process is cell differentiation^8,11,12,13,14^.

On the other hand, several studies of metamorphosis have been carried out using as experimental model salamanders from the *Ambystoma* genus. In this genus some species can not undergo metamorphosis and keep juvenile traits all their life cycle, in a phenomenon called neoteny, while others can metamorphose into terrestrial organisms^17,18,19^. In *Ambystoma mexicanum*, which is a neotenic species, the metamorphosis is induced by the application of exogenous Thyroxine (T4)^20^ and morphological changes were identified in different organs during metamorphosis. For example, in the metamorphic heart, similarities are observed in terms of cardiomyocyte dispersion and organization of the ventricular tissue^21^. However, the characterization at tissular level and the formation and organization of the valves and the atrial zone has not been thoroughly described. With respect to the lungs, an increase was observed in the alveolar cells that have a function in pulmonary respiration^21^. Changes were also found in the thyroid gland^22^, and in the size of the organism that decreases during metamorphosis^23^. Externally the epidermis of induced individuals is formed of keratinized stratified squamous epithelium^24,25^. Finally it was also found that genes involved in neural development are altered during metamorphosis^26^.

The previously described studies have been of value to understand in more depth the process of metamorphosis in urodele amphibians. However, it has been reprted for some neotenic species that T4-induced organisms do not develop normally and their viability is affected after the transformation, depending on the genetic background lines used^66^. Among the *Ambystoma* genus, several other species have the capacity to metamorphose in response to environmental changes, such as *Ambystoma velasci*, a facultative metamorphic species endemic to Mexico.

In this study we use *A. velasci* as the experimental model, in order to understand the transcriptional changes in respiratory and circulatory systems along the transition of the organism from aquatic to terrestrial environments. We selected heart, lungs and gills as the main target tissues to explore their transcriptional programs during five stages of metamorphosis. According to the gene ontology term-enrichment analysis, we identified the genes involved in synthesis, perception and response of TH in the three organs studied. In the metamorphic heart, we observed transcripts that are differentially-expressed involved in the differentiation of cardiomyocytes, particularly in the development of the interatrial zone, vasculogenesis and in the maturation of coronary vessels. In the lung, we observed genes related to hematopoiesis, angiogenesis and synthesis of the surfactant protein. In the gills, the most prominent biological processes identified are degradation of extracellular matrix, apoptosis and keratin production. This allows us to propose *A. velasci* as an excellent model to characterize the morphological and physiological changes caused by the diverse genetic programs underlying this interesting, and still intriguing, developmental process.

## Materials and Methods

### Sample preparation

Aquatic specimens of *Ambystoma velasci* of 13-15 cm of total length, were collected from Tecocomulco lake (Hidalgo, Mexico; 19.8649° N, 98.3910° W). These organisms were kept in glass containers with antichloride-treated water and weekly water renewal, with a 12/12 cycle of light/darkness in a constant temperature of 17°C. Organisms were all fed 5 times a week with a diet composed of red worm (*Lombricus terrestris)* and water flea (*Daphnia magna).*

### Metamorphosis induction and morphometric analyses

Metamorphosis was induced in 30 organisms with applications of 50 nM T4 hormone (Sigma T2376, St. Louis, MO) every third day^23,24,26^. During this process phenotypic changes that delimit one phase to the next occur. In this sense, a daily monitoring of the morphometric characteristics was performed measuring: Total length (TL), Snout-cloaca length (SCL), Total tail width (TTW), Tail dorsal width (TDW), Tail ventral width (TVW), Gill length (GL) and weight (g)^20,23^. We analyzed statistically the morphometric characteristics during the metamorphosis process, the metamorphosis stages were determined for TDW and GL characters (p<0.05). Using morphological traits, samples were collected in stage 0 (S0) premetamorphosis at 0 hours, stage I (SI) at 12 hours post-induction, stage II (SII) at 36 hours post-induction, stage III (SIII) or metamorphosis climax 6-12 days post-induction, and stage IV (SIV) or post-metamorphosis 17-23 days post-induction, respectively. To minimize biological variance, 15 individuals were sacrificed, using three individuals in each stage. Were anesthetized in 0.01% benzocaine and dissected the heart, lungs and gills. For the gills in SIV, a sample of skin tissue was sampled from the head region where the gills once were.

### Animal experimental procedures approval

The animal experimental procedures were performed according to the Mexican Official Norm (NOM-062-ZOO-1999) “Technical Specifications for the Care and Use of Laboratory Animals” based on the Guide for the Care and Use of Laboratory Animals “The Guide”, 2011, NRC, USA with the Federal Register Number # B00.02.01.01.01.0576/2019, awarded by the Secretaría de Agricultura y Desarrollo Rural (SADER) who is the federal authority that verifies the compliance of the Mexican Official Norm in Mexico. The Institutional Animal Care and Use Committee (IACUC) from the Center of Advanced Research and Advanced Studies (CINVESTAV) approved the project “Management and husbandry of *Ambystoma spp* and experimental processing of tissue for functional analyses and genetic expression” ID animal use protocol number: 0209-16.

### RNA extraction and Illumina sequencing

Total RNA was isolated using TRIzol® reagent (Invitrogen, Carlsbad, CA, USA), from 45 tissue samples (3 different tissues - Heart, Lung and Gills-) from 3 independent organisms for each of the 5 time points of metamorphosis tested. Each RNA sample was analyzed for quantity and purity with Bioanalyzer. From each tissue and time point, 3 independent RNAs were pooled and from such 15 pooled (3 organs/5 stages) RNAs, libraries were generated using TruSeq protocols and sequenced in 2×100bp format in Illumina HiSeq2500 platform. The sequencing was generated at the UGA-CINVESTAV genomics services.

### *De novo* transcriptome assembly and annotation

In order to obtain high quality readings, the readings obtained from the sequencing were filtered following a quality (greater than 20) using Trimommatic v0.36 software, and the sequences of the adapters were eliminated. The clean reads were assembled *de novo* using Trinity software v2.0.6^67^. After assembly, the libraries were combined (the contigs were put together in a single fasta file) and the similar sequences were grouped with CD-HIT-EST, which align the sequences in pairs and group them if they have an identity> 95% and coverage> 95%, already with the groups generated, only the largest contig was reported, eliminating the redundancy of the libraries and leaving only the transcripts that best represent the transcriptome. The high quality readings were mapped to the assembled transcriptome with Kallisto v0.42^68^ to obtain an estimate of the relative levels of expression in each organ of five stages in metamorphosis. Each sample was quantified separately reporting the observed and adjusted counts of each sequence, in addition to its value in Million Count (CPM). The data for each sample were described in a table, which reports for each representative transcript sequence its value in effective (adjusted) counts, that is the entry point of edgeR.

### Differential transcripts expression analysis

In edgeR, the data in estimated counts was normalized (the default algorithm is the TMM) and the expression tables were generated. The selection of differential transcripts was made, comparing the conditions against stage 0 or control, selecting transcripts with Fold Change (enrichment)> 1X and False Discovery Rate (FDR)<0.05. From this list of differentially expressed transcripts, we considered three selection criteria for the transcripts of interest; a). The values of *p-value* and FDR, b). The expression patterns, the transcripts at each stage of the metamorphosis were also considered, that is, those that will only be expressed at a specific stage, gradually increasing, gradually decreasing and those that remain unchanged in their expression, c). Gene Ontology analysis, the biological event in which these genes were involved.

### GO enrichment analysis

Subsequently, an enrichment analysis of Gene Ontology (GO) categories were performed for the genes that were found as differentially expressed and enrichment in each organ-stage, in PANTHER Classification System software^69^. GO is a bioinformatics tool that unifies the criteria for gene annotation and the biological functions attributed to them. Panther GO, database covers five domains: Molecular Function MF (nine subcategories), Biological Process BP (15 subcategories), Cellular Component CC (eight subcategories), Protein Class PC (26 subcategories) and Pathway (162 subcategories). Only GO categories with a FDR<0.05 were considered.

### qRT-PCR

To determine the expression patterns of specific DEGs, we selected a group of genes related to the pathway of THs suchs as those for thyroid hormone synthesis (*Avtg* and *Avdio2*), thyroid hormone receptors (perception) (*Av*TRα, *Av*TRβ, *Av*RXRα and *Av*RXRβ) and thyroid hormone response (*Avhbe* and *Avhba)* for the three organs. As well as for DEGs which homologs have been related to the biological processes of heart formation and function. The transcripts of these genes were quantified in each organ and stage by qRT-PCR. For this assay, a different group of organisms of those used for the RNAseq approach were used. Total RNA was isolated, using TRIzol® reagent (Invitrogen, Carlsbad, CA, USA), from 3 different tissues (Heart, Lung and Gills) from 3 independent organisms for each of the 5 time points of metamorphosis tested (in total 45 RNA samples). Each RNA was converted independently in cDNA using SuperScript II Reverse Transcriptase (Thermo Fisher Scientific). Real-time quantification was performed using SYBR→ Green Master Mix in 10-μL reactions. The primers sequences used for qRT-PCR amplification are listed in STable 12. PCR was performed using a program of 95°C for 10 min and 40 cycles of 95°C for 15 sec and 60°C for 1 min (STable. 12). The expression levels of genes were normalized to the expression of GAPDH. The standard curves were quantified PCR products of 10-fold serially diluted, and linear regression equations and R^2^ was made in Microsoft Excel (STable. 13). The real-time PCR amplification efficiency was calculated as E=[10^(−1/slope)^]-1. The relative expression levels of genes were calculated as 2^-⊗⊗CT^. Technical triplicate was done for each primer pair tested. The statistical analyzes were performed on the mean and standard deviation for the technical triplicates of each biological replicate. (For more details consult the KRT table -Key Resources Table-).

## Results

### Characterization of the metamorphosis stages in *Ambystoma velasci*

The metamorphosis was induced with 50 nM of T4 and the periodicity of T4 application is shown in Figure 1 (a). To define the major stages of metamorphosis in *A. velasci*, we recorded data of diverse morphometric criteria every day until the end of the process (Fig. 1b). Our morphometric analyses are quite similar to those previously reported in several other studies for *Ambystoma* species metamorphosis^20,25^, the parameters measured were total length (TL), snout-cloaca length (SCL), total tail width (TTW), tail dorsal width (TDW), tail ventral width (TVW), and gill length (GL). Our statistical analysis showed that the measurements that constantly reduce during the metamorphosis process are: GL (*p*<0.05) and TDW (*p*<0.05) (Fig. 1c, SFig. 1, and STable 1). At SI and SII, a tendency was observed in the reduction of 14-24% of the GL with respect to the initial length. In SII, the weight (STable 1), TL and SCL remained stable. At SIII, GL continues to decrease their length up to 50-75%, TDW is reduced between 20-40%. By SIV, branchial tissue has fully degraded and TDW is reduced between 85-100% in this stage (Fig. 1c).

**Figure 1.**
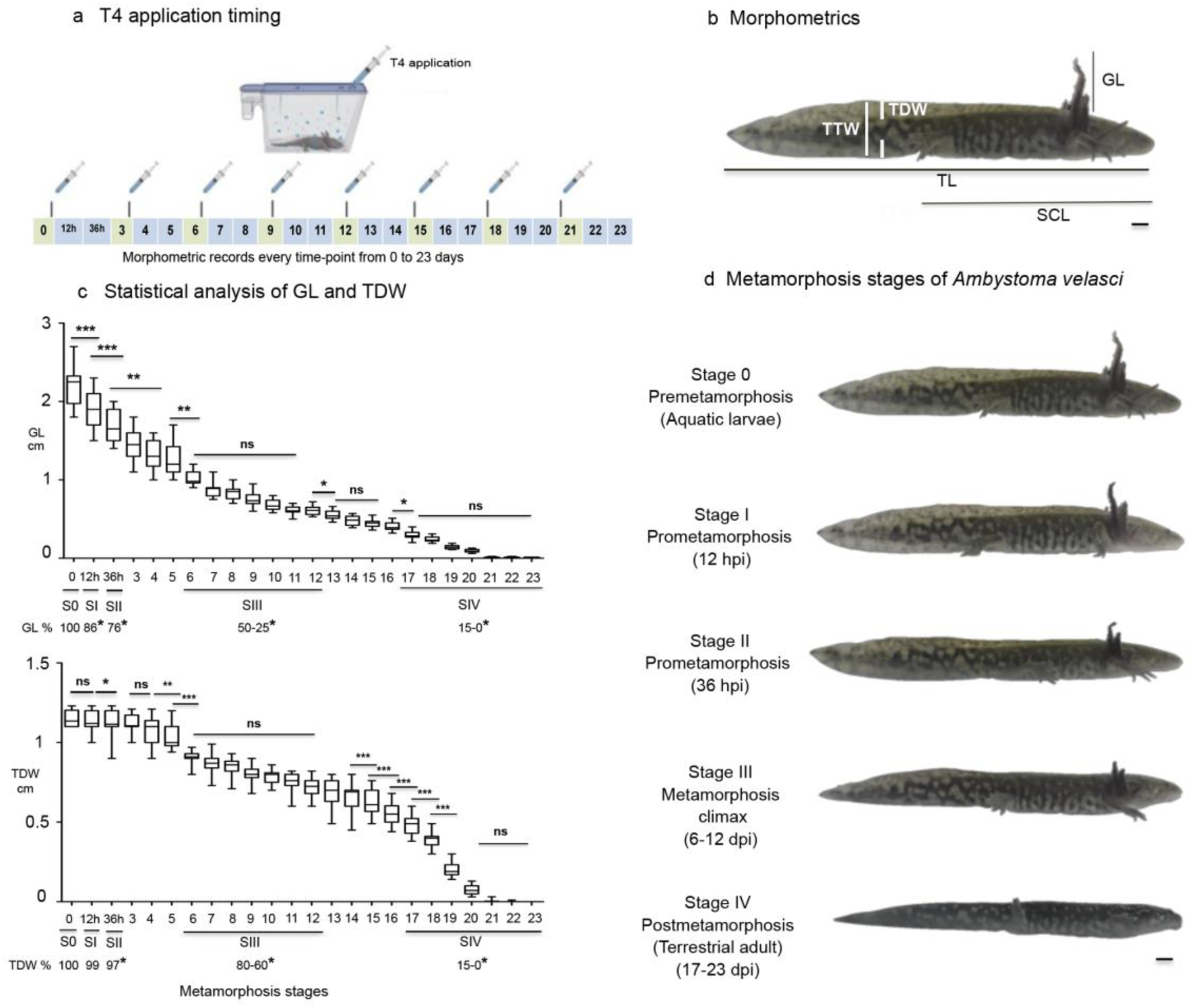
Determination of metamorphosis stages in *A. velasci* organisms. (**a**) Time points for application of 50 nM of T4 and for morphometrics recording from day 0 to day 23 postinduction. (**b**) The morphological parameters measured were total length (TL), snout-cloaca length (SCL), total tail width (TTW), tail dorsal width (TDW), tail ventral width (TVW), and gill length (GL). The stages of metamorphosis were established based on (**c**) statistical analysis of the morphometric characteristics which allowed us to divide the process in (**d**) stage 0 (S0) where the juvenile organisms are aquatic, stage I (SI) or prometamorphosis at 12 hours postinduction (hpi), in which a reduction of 14% of the GL was observed, SII at 36 hpi, where GL has reduced by up to 24%, stage III (SIII) or Climax a 6-12 days postinduction (dpi) is characterized by the reduction of 50-75% of the gills and stage IV (SIV) a 17-23 dpi when the gills have completely disappeared and the organism has fully transformed to terrestrial (n=30), the X axis the days and metamorphic stages, and Y the morphometrics characteristics. Scale bar is 1cm. For a more detailed view of the morphological changes see Supplementary video 1, Metamorphosis stages of *A. velasci*. Illustration in (**a**) created with Biorender.com.

Based on the changes measured for GL and TDW, we classified the *A. velasci* metamorphosis process in the following stages: Stage 0 (S0) or premetamorphosis, Stage I (SI) at 12 hours post-induction (hpi), Stage II (SII) at 36 hpi, Stage III (SIII) metamorphosis climax a 6-12 days postinduction (dpi), and Stage IV (SIV) postmetamorphosis a 17-23 dpi (Fig. 1d).

### *De novo* transcriptome assembly and annotation

From the previously described characterization, we selected five metamorphosis stages (S0 to SIV) in *A. velasci* to profile and analyze the transcriptional landscape of heart, lungs and gills during metamorphosis. For each stage, the heart, lungs and gills of 3 organisms were dissected and total RNA extracted. The concentration and integrity of the RNA was verified and a pool made from the 3, independently isolated, RNAs of each organ and stage was used for the preparation of 15 libraries. Paired end sequencing delivered 58,191,270 raw reads for heart, 60,077,384 for lung and 66,903,152 for gills, these reads represent the total read combining the five stages sampled per organ. Processed and clean reads were assembled in a total of 187,627 coding transcripts for heart, 211,100 coding transcripts for lungs and 226,738 coding transcripts for gills, the global and the stage-specific transcripts and reads are resumed in Table 1. In addition, we verified the reads coverage per library to the full assembled transcriptome, we found that >93% raw reads aligned to the transcriptome in each library (STable 2).

**Table 1.** Global transcriptional analysis of diverse organs in five metamorphosis stages. Numbers resulting from diverse analyses after sequencing-assembly and coding transcripts in heart (H), lung (L) and gills (G), in five different metamorphosis stages of *A. velasci*, where S0 represents the premetamorphic organism, SI at 12 hours postinduction (hpi) with T4, SII at 36 hpi, SIII at 12 days postinduction (dpi) and SIV at 23 dpi.

| Metamorphic stage per organ | Number of reads | Number of transcripts | Number of coding transcripts |
| --- | --- | --- | --- |
| **Heart-S0** | 8,305,940 | 56,278 | 31,516 |
| H-SI | 8,151,630 | 61,837 | 35,510 |
| H-SII | 12,083,466 | 66,547 | 40,128 |
| H-SIII | 13,806,442 | 85,944 | 45,388 |
| H-SIV | 15,843,792 | 60,768 | 35,085 |
| Total | 58,191,270 | 331,374 | 187,627 |
| **Lung-S0** | 13,637,330 | 87,694 | 44,163 |
| L-SI | 14,688,672 | 89,706 | 46,052 |
| L-SII | 13,030,808 | 93,672 | 47,015 |
| L-SIII | 10,829,318 | 84,116 | 43,679 |
| L-SIV | 7,891,256 | 47,726 | 30,191 |
| Total | 60,077,384 | 402,914 | 211,100 |
| **Gills-S0** | 12,575,638 | 77,054 | 40,264 |
| G-SI | 13,492,506 | 83,124 | 43,216 |
| G-SII | 13,615,934 | 84,168 | 42,595 |
| G-SIII | 15,640,802 | 86,286 | 46,068 |
| G-SIV | 11,578,272 | 99,674 | 54,595 |
| Total | 66,903,152 | 430,306 | 226,738 |

### Differential Genes expression (DGE) analysis per stage and organ during *A. velasci* metamorphosis

In order to identify differentially expressed genes (DEGs) in each stage and organ, we performed comparisons among heart, lung and gills and between stages per organ, using S0 as the baseline (S0 *vs*. SI, SII, SIII and SIV) with the edgeR software. Interestingly in heart the number of DEGs is higher at SIII compared to lung and gills, where an increase can be observed clearly at SIV (Fig. 2).

**Figure 2.**
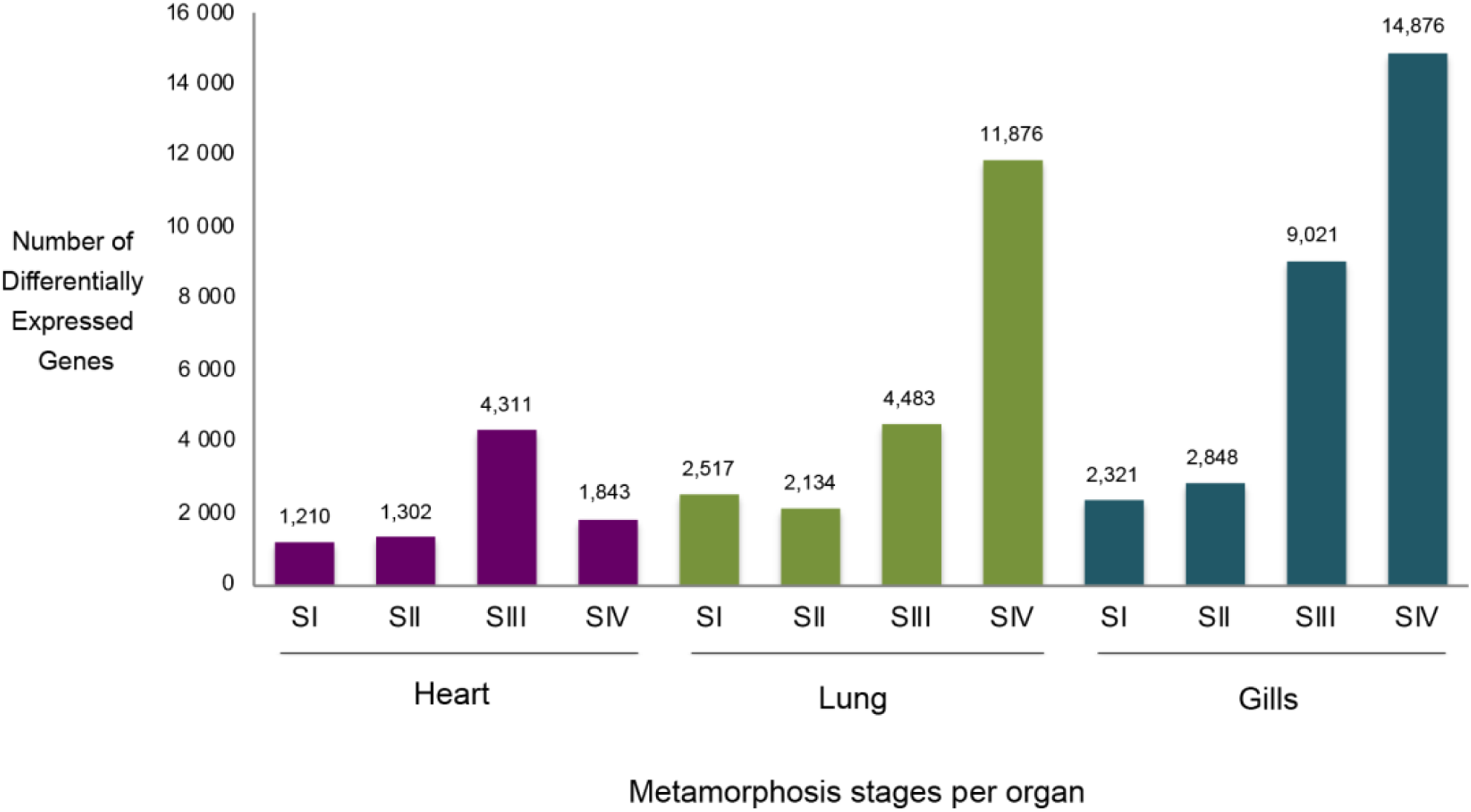
Differential Gene Expression (DGE) in diverse organs during the metamorphosis stages. In the X axis the 3 different organs at each metamorphosis stage are labeled. Purple bars for the heart, green for lung and turquoise for gills. The S0 represents the aquatic organisms, SI-12 hours postinduction (hpi) with T4, SII-36 hpi, SIII-12 days postinduction (dpi) and SIV-23 dpi. The Y axis represents the number of differentially expressed transcripts. Stage 0 is used as a baseline for comparison.

### Stage /organ-specific Up and Down regulated transcripts

The identified DEGs among the five stages for each *A. velasci* organ analyzed were clustered and represented in heat maps (Fig. 3a-c). We divided the DEGs patterns into eight different groups according to their Fold Change (enrichment)>1X, *p-value*<0.05 and an adjusted *p-value* for FDR<0.05 (Fig. 3d) for each organ. Such clusterization allowed us to identify stage-specific and highly expressed genes, enriched genes along all the stages, genes which expression gradually increase during the process and those gradually decreasing (Fig. 3d).

**Figure 3.**
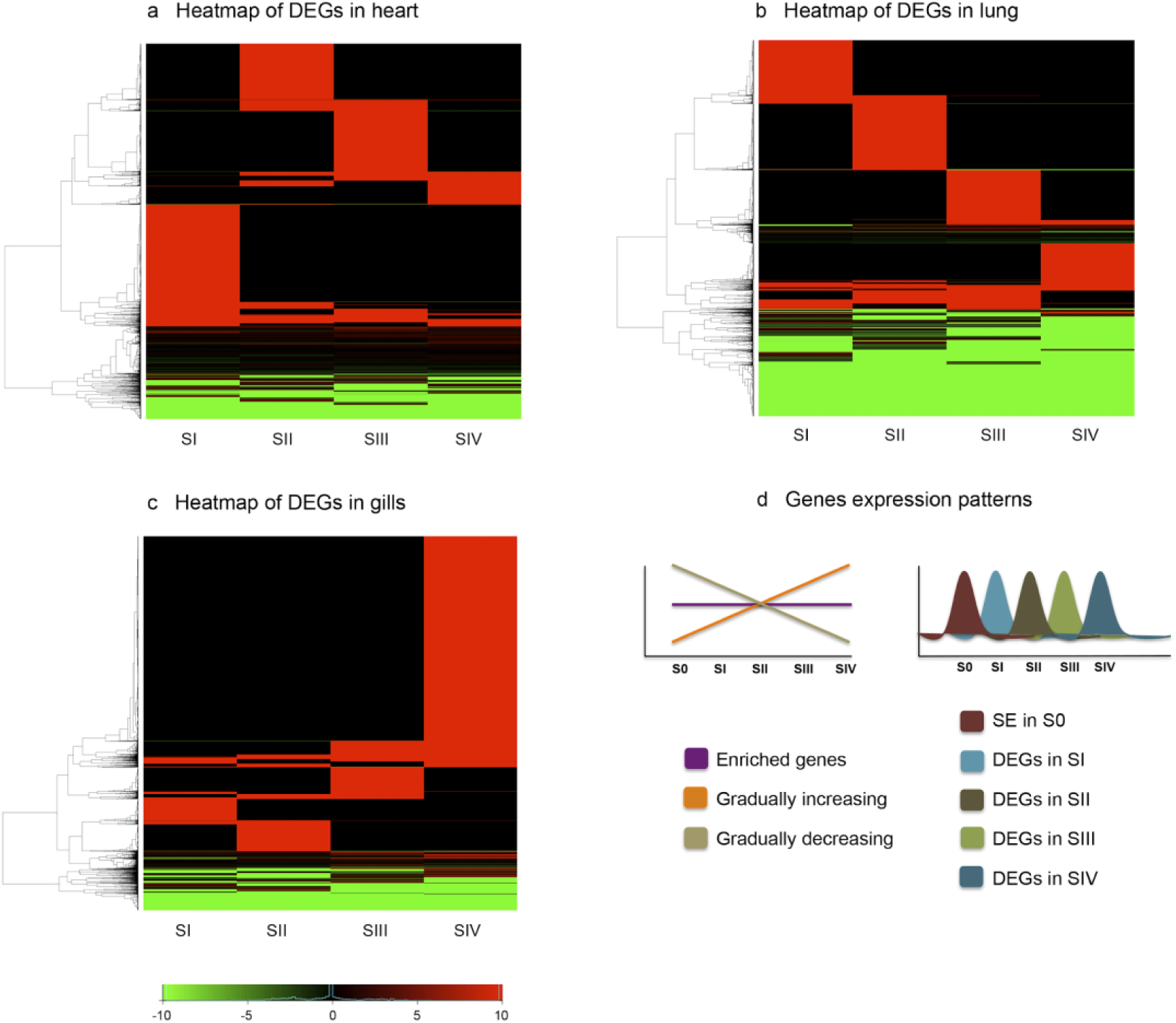
DEGs per organ and metamorphosis stage. Heat maps showing patterns of differentially expressed genes in heart (**a**), lung (**b**), and gills (**c**), in each of the metamorphosis stages analyzed, S0, SI-12 hpi, SII-36 hpi, SIII-12 dpi and SIV-23 dpi. Stage 0 was used as the reference stage for DGEs analyses. (**d**) Graphic representation of all the patterns followed by differentially expressed genes. Genes were grouped by expression pattern as enriched or stage specific (S0, SI-12 hpi, SII-36 hpi, SIII-12 dpi and SIV-23 dpi). Besides stage-specific enriched pattern, those with enriched expression during metamorphosis, those with increasing gradient expression and those with gradually decreasing expression were identified and grouped. SE in (**d**) for S0 means Stage-enriched.

### Gene ontology analysis of DEGs per stage and organ

To shed light on the putative molecular pathways that underlie the biological processes along metamorphosis, the 4,064 DEGs in heart, 4,107 in lung and 8,265 in gills, were subclassified into eight major clusters: Enriched genes, gradually increasing, gradually decreasing, stage-specific genes in S0, SI, SII, SIII, and SIV (STable 3-5, Figure 3). The genes grouped in each of the eight different clusters for heart, lung and gills were then analyzed to define enriched functional categories by using PANTHER 14.0 Classification System (STable 6-8 GO categories and STable 9-11 GO subcategories).

The GO analysis reveals that different biological processes are involved in the organs of the respiratory and circulatory systems during metamorphosis. Globally, in the enriched-genes cluster in all tissues the most represented subcategories are binding (GO:0005488), catalytic activity (GO:0003824), and developmental process (GO:0032502). The most enriched Biological Process (BP) subcategory in heart and gills was metabolic process (GO:0008152), while for lung it was cellular process (GO:0009987) (Fig. 4b, SFig. 2 and 3). Interestingly in the subcategory metabolic process (GO:0008152), for the three organs, we identified the homologous of the *hbe1* gene, which codes for a type of hemoglobin that is expressed in embryos and that has been described as a TH-responsive gene^33^.

**Figure 4.**
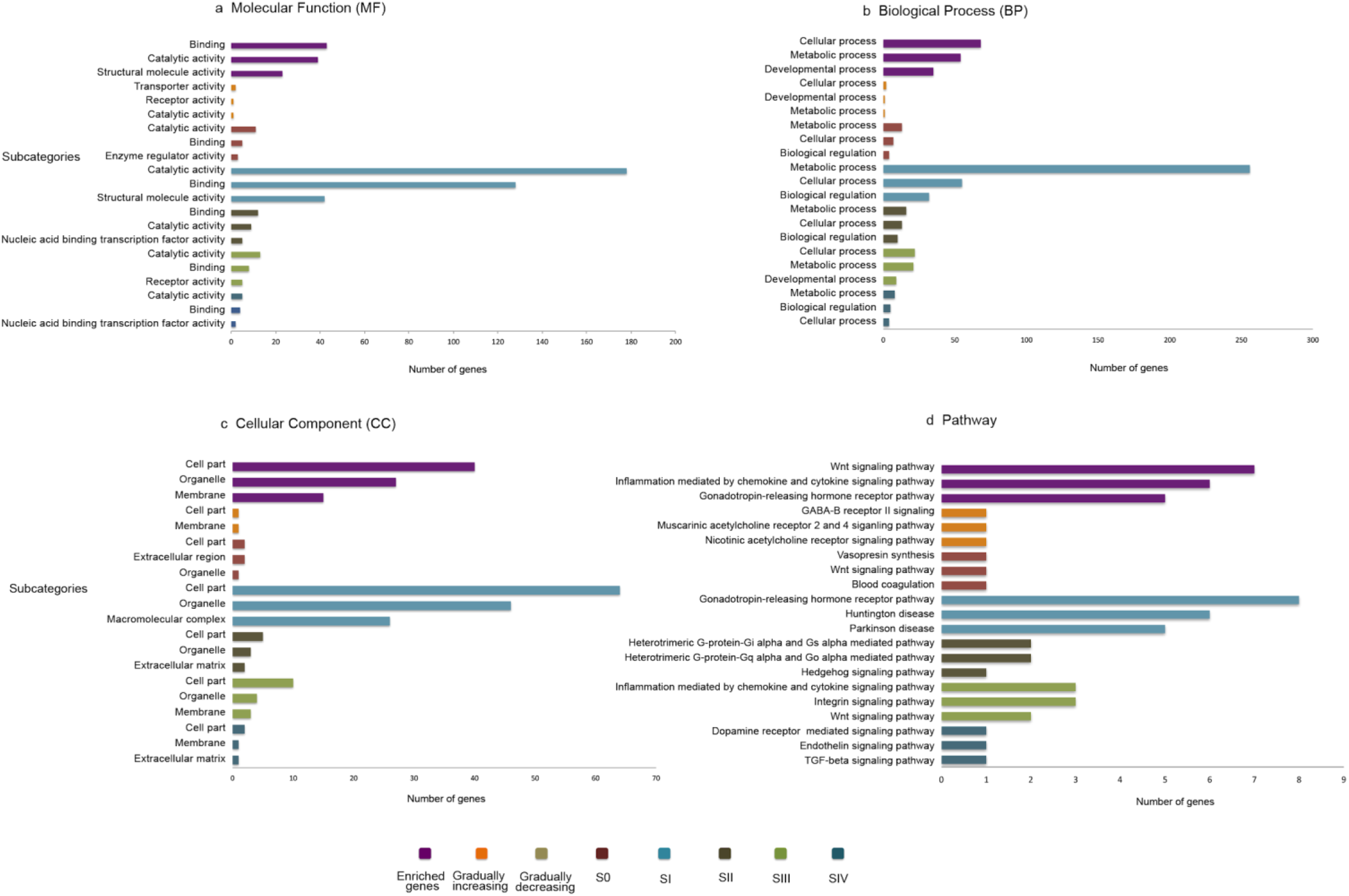
Gene Ontology analysis in heart during metamorphosis. Enriched functional categories of DEGs in heart by expression pattern during metamorphosis. Stage 0 is used as reference. The major categories analysed were (**a**) Molecular Function category (MF), (**b**) Biological Process (BP), (**c**) Cellular Component (CC) and (**d**) Pathway. In the Y axis the subcategories of the gene ontology are represented, in the X axis the number of genes enriched in each subcategory. The clusters are represented by different colors: purple for heart-enriched genes compared to lung and gills, orange for gradually increased expression, red for transcripts enriched in S0, blue for SI-12 hpi, brown for SII-36 hpi, green for SIII-12 dpi and turquoise for SIV-23 dpi.

In heart-enriched genes, as part of the binding subcategory (GO:0005488), we identified genes that are involved in the differentiation of cardiomyocytes (*gata4*^27^), and the heart atrial zone development (*myl7*^28^). Also genes that participate in heart morphogenesis such as *myh4*, MYH4 is considered a structural constituent of the cardiac muscle^35^. We also identified *bmp4* that codes for BMP4, a ligand involved in the TGF-β pathway which is expressed in developing and adult heart^29^. Another gene differentially expressed in this subcategory is *nkx2.5*, which is expressed during embryogenesis and adulthood in human heart^32^.

Interestingly, in the cluster of genes enriched in the lung (STable 7 and 10, SFig. 2a), one of the most represented subcategories is binding (GO:0005488), that is related to lung development. In this subcategory we identify the homologous gene for Tbx4^41^ and Foxa2. Foxa2 is involved in the synthesis of surfactant protein that is carried out in the alveolar cells type II in mammals^62^. While some genes enriched in gills, as part of binding (GO:0005488) subcategory (STable 8 and 11, SFig. 3), such as *pkp1* and *lgals1*, are involved in ectoderm morphogenesis during embryonic development in mammals^49^ and in regulating apoptosis, cell proliferation and differentiation^50^.

Focusing on the GOs for the gradually decreasing transcripts, we observed that in the lung, the *Avhoxc8* expression is decreasing during metamorphosis, correlating with the functions of this gene in cell proliferation and differentiation^42^. With a similar expression pattern, we identify *rhcg* in the gills, that codes for Ammonium transporter type C, a membrane protein that excretes ammonium and plays a role in pH regulation in aquatic animals^51^.

Interestingly in SI in the heart, we identified as the most enriched subcategories catalytic activity (GO:0003824) and binding (GO:0005488). In these subcategories, we found *tgfbr1* and *smad4*, these both are factors of the TGF-β pathway. This pathway is critical for maintaining cardiac function and for cardiomyocyte survival, and has been described as activated in response to THs^29^. Another biological process identified in the heart is that of the homeostasis of the organs^36^ in which *stx5* is upregulated in this process. Regarding the lung, we were able to identify the gene *jag2*, Jagged2 is a ligand than binds NOTCH1, with important roles in embryo development and stem cells maintenance^43^. Another enriched subcategory is catalytic activity (GO:0003824), which includes *uts2r*, a gene coding for the Urotensin II receptor, which is highly expressed in the human lung and has been implicated in regulating the respiratory physiology in mammals^44^. Particularly in the gills we identified *hoxa7*, involved in regulating the expression of differentiation-specific keratinocyte genes^52^. As results of the analysis to identify enriched functional categories of BPs in the heart at SII, we found that the subcategory developmental processes (GO:0032502) is enriched, in which the expression of genes related to the development of the vasculature of the coronary arteries and the atrial septum formation is high, such as *gli3* and *pka* that participate in the SHH pathway^34^. As well as *myh6*, that has been reported as expressed in the developing heart^56^. We also identify *adora3* that codes for the Adenosin receptor 3, that participates in the regulation of heart rate^37^. In the lung, some upregulated genes involved in hematopoiesis and cilia structure and function in the respiratory system, as *mecom*^45^ and *dnah11*^46^ respectively. We also identified in the gills the subcategory metabolic process (GO:0008152) in which we found *krt7*, Keratin type II involved in development, differentiation and maintenance of simple and a transitional epithelia^53^, a gene which expression has been shown to be regulated by *hoxa7*.

Interestingly, in the metamorphosis climax or SIII, in the heart and lung we identified in the binding subcategory (GO:0005488) the up-regulation of genes which are related to the immune response. In the heart, *hla* is expressed in the microvascular endothelium in endocardial cells from humans with myocarditis^30^. Some of the genes involved in the development of the vasculature of the coronary arteries and the atrial septum formation, are also upregulated at SIII. We also observed an enriched expression of *gata4*, that in addition to being enriched in the heart is involved in the differentiation of cardiomyocytes. Another of the processes that are enriched is that of homeostasis of the heart, we identified as upregulated *Avmas*, which human homologue participates in the Renin-Angiotensin-Aldosterone (RAAS) pathway, that modulates liver, kidney and heart homeostasis^57^. While in the lung we found upregulated *Avfcn1*, which homologue in human codes for Ficolin-1, a protein involved in complement system activation of the immune system^47^. In the same subcategory in gills we find *Avadamts10*, ADAMTS10 is a metalloproteinase that in humans, that participates in morphogenesis, remodeling, and vascularization^54^.

When the metamorphosis reaches the end at SIV, organisms are capable of living in a terrestrial environment. In such a stage we identified among the upregulated genes in the BP category in the heart, *Avthsd7.* In humans, this gene codes for Thrombospondin 1, a component of the ECM (extracellular matrix) and is involved in maintaining heart structure and blood vessels, the control of cell growth, regulation of diastolic stiffness, as well as in tissue repair when cardiovascular damage and inflammation occurs^31^.

One of the last processes of pulmonary maturation in mammals involves the formation of alveoli, which are specialized structures in gas exchange^48^. In this sense in the lung in the subcategory binding (GO:0005488), we identified upregulated *angptl3* that codes for Angiopoietin 5, which is involved in the processes of angiogenesis and alveolarization^48^. Also in this stage the gill tissue has disappeared completely (Fig. 1d), interestingly we find *timp2*, that is a metalloproteinase inhibitor (TIMP2)^54^.

We analysed the functional subcategories enriched using as a filter of analysis the Pathway category in the heart (Fig. 4d), we found pathways related to myocardial function. In the cluster of gradually increasing genes in the GABA-B receptor II signaling subcategory (P05731) we identify *Kcnj3* that in mammals is involved in the depolarization and repolarization of the cardiac action potential^38^. In the SI the most enriched subcategory was Integrin signalling pathway (P00034), we found genes related to cell proliferation and chamber maturation in cardiac development in mammals^39^. In the SIII we also identified genes involved in the formation of the coronary vessels in the atrial-ventricular zone in mouse^40^, grouped in the Inflammation subcategory (P00031). Regarding the lung, the pathway most represented in the process of metamorphosis is the Angiogenesis pathway (P00005).

### Expression patterns validation for diverse *A. velasci* homologs of TH-related genes

In order to determine the expression patterns of genes whose homologs have been shown in other organisms to be involved in the TH pathway, we analyzed by qRT-PCR their transcript levels along the diverse stages of metamorphosis in heart (Fig. 5), as well as in lungs and gills (SFig. 4 and 5). The expression patterns of the four thyroid hormone receptors *Av*TRα, *Av*TRβ, *Av*RXRα and *Av*RXRβ were validated by qRT-PCR (Fig. 5a). Higher levels of the transcripts corresponding to *Av*TRα, *Av*RXRα and *Av*RXRβ receptors were observed during SIII, while the transcript levels of *Av*TRβ receptor are, almost exclusively, expressed in SIV. Our qRT-PCR-based analyses also show that by stages SIII and SIV the transcript levels of *Avtg*, the homolog of the protein involved in the synthesis of the major precursor of the thyroid hormone^55^, are induced. We also confirmed that higher levels of transcripts of *Avdio2* (Fig. 5b), were present in SIII. The transcriptional induction of the thyroid hormone-responsive genes *Avhbe* and *Avhba*^26^, which expression is higher in SI and SII, respectively, was also confirmed by qRT-PCR (Fig. 5c).

**Figure 5.**
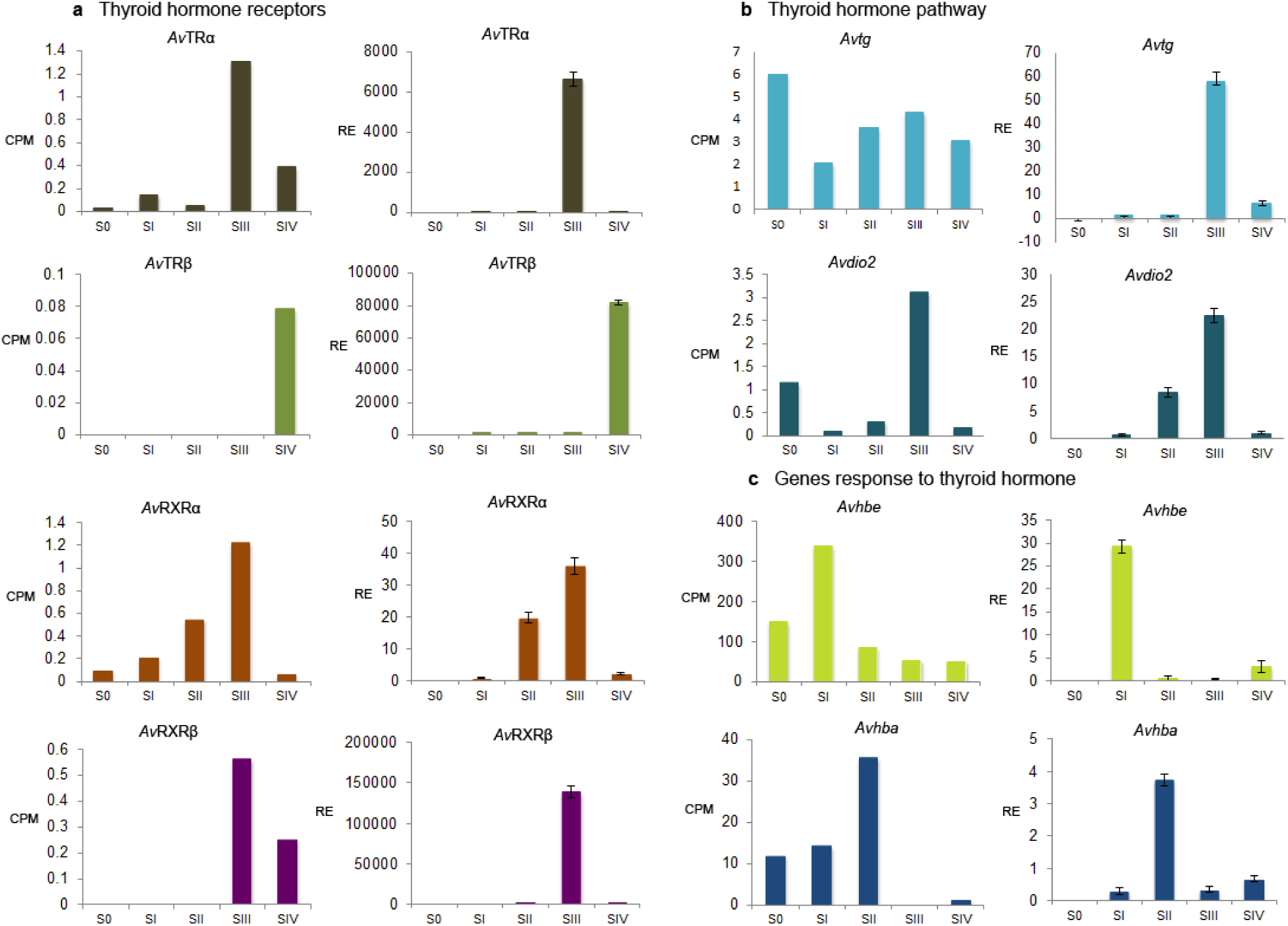
Validation of DGE patterns of *A. velasci* homologs for genes involved in the thyroid hormone pathway in the heart. The X axis represents the metamorphosis stages S0, SI-12 hpi, SII-36 hpi, SIII-12 dpi and SIV-23 dpi. (**a**) Expression patterns of *A. velasci* homologous genes for thyroid hormone receptors: *Av*TRα, *Av*TRβ, *Av*RXRα and *Av*RXRβ (**b**) Genes involved in thyroid hormone synthesis, *Avtg* and *Avdio2*. (**c**) Genes that have been reported as responsive to thyroid hormone; *Avhbe* and *Avhba*. The counts per million (CPM) value represents the transcriptome result. The relative expression (RE) is the result of qRT-PCR analysis, the columns and bars represent the mean and the standard deviation of the three biological replicates with three technical replicates each individual samples.

### Validation of DEGs of diverse *A. velasci* homologous genes involved in heart development and function

We also selected a group of genes involved in heart development, to determine their expression patterns by quantifying their transcript levels using qRT-PCR. According to our transcriptomic data, in the gradually increasing genes (Fig. 3d) was *Avsmad6*, our qRT-PCR analyses showed that the transcript levels of this gene increase through all metamorphosis stages (Fig. 6). In contrast, the gradually decreasing expression pattern observed for *Avmyl7* in the transcriptomic analysis, was not confirmed by qRT-PCR, instead a very specific expression in SI was observed (Fig. 6). The SII-enriched expression of *Avmyh6* was confirmed. The expression patterns of *Avtbx1* and *Avgata4*, which are mostly expressed in SIII, were also validated. In cluster SIII we also predicted and confirmed as highly expressed *Avgli3, Avpka, Avtbx1* and *Avmas* genes (Fig. 6).

**Figure 6.**
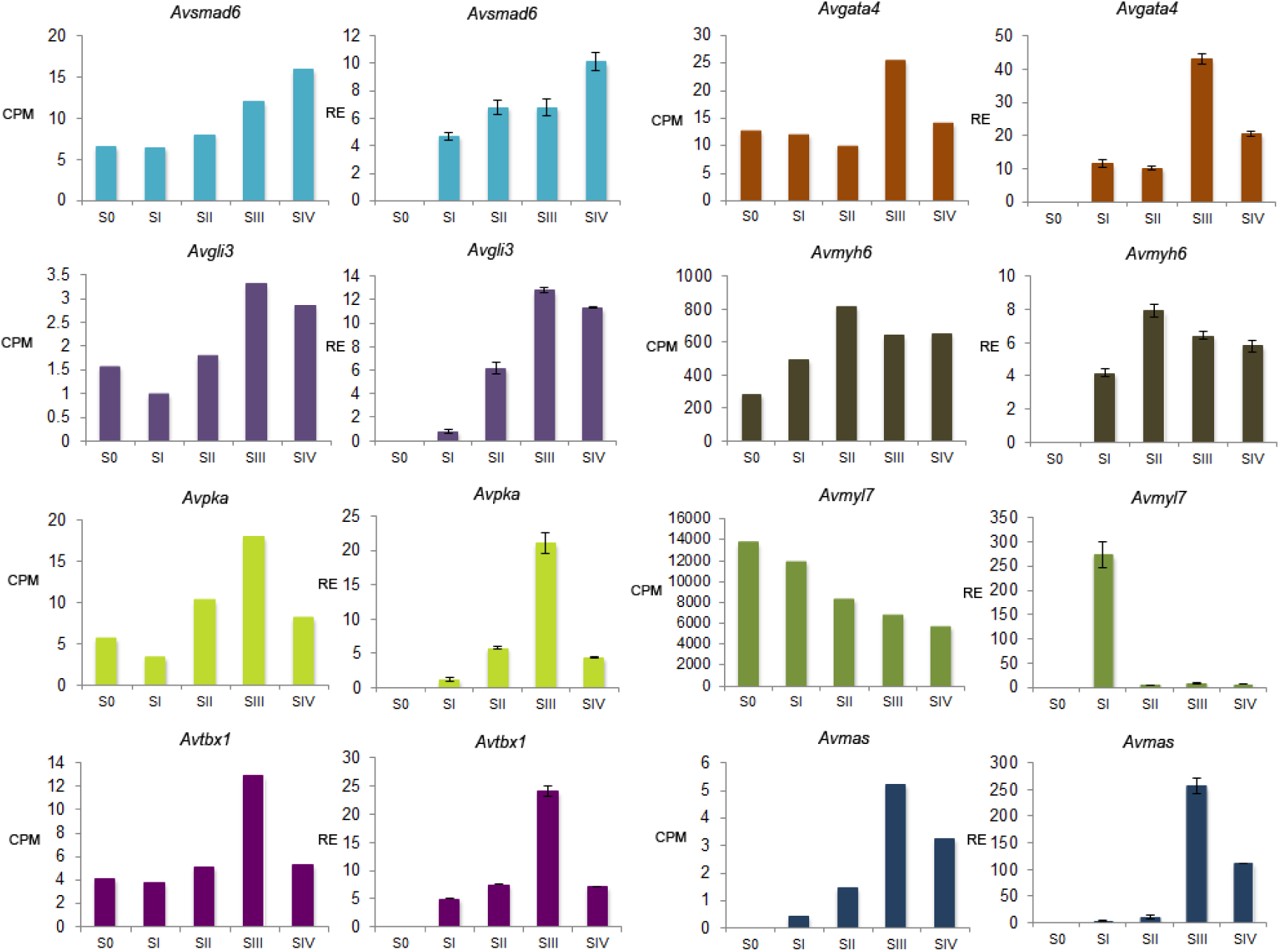
Validation of DGE patterns of transcripts involved in heart development and function. For each gene the transcriptomic profile is shown by the CPM value and the RE obtained by qRT-PCR analysis. The columns and bars represent the mean and the standard deviation of the three individual samples. The X axis represents the metamorphosis stages S0, SI-12 hpi, SII-36 hpi, SIII-12 dpi and SIV-23 dpi. The expression patterns of *A. velasci* homologous genes were grouped by their stage-specific expression pattern in heart during metamorphosis. These transcripts set code for proteins involved in heart development.

## Discussion

In this study, we report the transcriptomic profiling during the metamorphosis process of *A. velasci*, for heart, lungs and gills. *A. velasci* is a facultative metamorphic organism endemic to Mexico, which becomes terrestrial in function of the environmental conditions. In contrast with neotenic species, terrestrial *A. velasci* salamanders are fully viable^66^. Our work represents the first one describing the transcriptional landscape of circulatory and respiratory organs in a metamorphic facultative species of the *Ambystoma* genus.

We first focused on the analysis of the transcriptional behavior of genes related to the THs pathway. In the case of thyroglobulin and *dio2* we observed an increase in expression in SIII. This result is similar to that described in *Microhyla fissipes*^4^ and coherent if we consider that THs are synthesized from thyroglobulin. The regulatory function of THs is mediated by thyroid hormone receptors^4^. We found four transcripts in *A. velasci* which seem to be homologous of TRα,TRβ, RXRα and RXRβ. The expression levels of *AvTRβ* are lower in the three organs compared to those of *AvTRα*, which is consistent with the observed behavior for these genes in the metamorphosis of other amphibians^6^. An increase in transcript levels of *AvTRα* is observed during SIII, this is similar to levels in metamorphosis climax in *Xenopus* and *Rana catesbeiana*^*6*^. It was demonstrated in mammals that the heart *TRα* expression is essential to maintain cardiac function after birth^63^. *Av*TRβ is upregulated in SIV in all three organs during metamorphosis, this result is in contrast to that described for *Xenopus*, which describes an initial increase in the ortholog gene during metamorphosis^6^. This result suggests that *AvTRα* transcriptionally responds initially and maintains the metamorphosis process until the climax in *A. velasci*.

Regarding the TH-responsive genes, we identified that the transcripts of hemoglobins *hbe* and *hba*^33^, are upregulated in heart, lungs and gills. *Avhbe* transcripts are more abundant in SI, while those of *Avhba* increase in SII similarly to what occurs during *Xenopus* metamorphosis^6^. Our results also suggest a similarity between the expression patterns of *Avhbe* and *Avhba* in SI and SII during metamorphosis in *A. velasci* and those of hemoglobins E and A during embryogenesis and in perinatal life of humans and mouse^33^. Since Hemoglobin A has a better affinity for oxygen molecules^33^, we speculate that *Avhba* expression increases during the transition from an aquatic to a terrestrial environment to cope with the drastic change in oxygen availability.

In SI at 12 hpi, we identify genes for THs response expressed in the heart, as well as involved in TGFβ pathway such as *Avtgfbr1, Avsmad4, and Avsmad6*^29^. The TGFβ pathway plays a direct role in heart development and has been associated with several pathogenesis related to heart failure^29,58^.

In SII, we identified groups of genes expressed in heart that in vertebrates are related to the formation of coronary vasculature (*gli3* and *pka*^34^), cardiomyocyte differentiation (*myh6*^56,58,63^) and atrial septation (*gata4*^27,60^). Some of these genes remain highly expressed in SIII. The formation of cardiac septa in vertebrates is carried out in different stages of morphogenesis. In *Mus musculus*, it occurs between stages E8.0-E14.0, while in *X. laevis* it occurs before metamorphosis in stages NF41-NF46. In *A. mexicanum*, the interatrial septum is formed during postembryonic development at stage 57 (subadult), but there is no evidence of valve formation between both chambers^67^, on the other hand only changes have been observed in the number and organization of cardiomyocytes in the ventricular zone, in such study authors focused in the ventricular zone and not in the atrial zone^21^. However, in metamorphic organisms such as *A. tigrinum*, after metamorphosis the main perfusion pathway is down to the entire length of the pulmonary artery that is located in the atrial zone^21^. The transcriptional profiles of the genes related to the morphogenesis of the atrial septum and the coronary vasculature suggest that this process begins during the SII of metamorphosis in *A. velasci*.

During lung SII, we identified genes associated with the stimulation for functional maturation of the surfactant-producing alveolar cells^58^. Such changes correlate with the gradual increase of *Avfoxa2* transcripts observed in lungs transcriptomes in our study. In mice, Foxa2 protein plays a role in lubricating the lung with the production of surfactant protein preventing collapsing^62^. This suggests that the lung bud that is present in aquatic organisms starts to reprogram transcriptionally in order to improve its gas exchange capacity to adapt to the terrestrial environment. In mice, increased levels of available THs coincide with the acceleration of alveolar septation, suggesting that THs enhances the structural development of lungs^58^. This developmental process involves *mecom*^45^ which regulates hematopoiesis and *dnah11*^*46*^ that regulates the formation of the cilia structure in the respiratory system. In our study, the transcripts of *Avmecom* and *Avdnah11* are enriched in SII, which suggests that the pulmonary bud present in aquatic organisms is being differentiated during metamorphosis until it becomes a terrestrial functional lung.

At the metamorphosis climax (SIII) of heart, the transcript levels of *Avtbx1* are upregulated. It has been shown in *M. musculus* that the upregulation of this gene is associated with the formation of the aorto-pulmonary septum, which divides the aorta from the main pulmonary artery during embryogenesis (E9.0-E11.0). The pulmonary artery develops before birth when the pulmonary circulation is required for blood oxygenation^61^. This allows us to suggest that during the SIII of the metamorphosis in *A. velasci* a process similar to the development of the pulmonary vein is carried out, a phenomenon described in mammals in embryonic stages, which could be linked to the change of aquatic habits to terrestrial ones. We also identified *Avitga4* as an upregulated transcript. In mammals, ITGA4 is involved in the formation of the coronary vessels in atrial-ventricular zone during embryogenesis^40^. The transcript levels of the *Avmas* ortholog are upregulated in heart. In humans, MAS functions as a receptor involved in the RAAS pathway which modulates liver, kidney and heart homeostasis^57^.

In SIV we identified genes that are putatively involved in alveolarization, a major process during late lung development. In such a process the formation of alveoli, the main gas exchange units, occurs in a THs-dependent manner^58^. In mammals the process of alveolarization starts after birth and hypothyroid lungs show progressively larger spaces, with less alveolar septations resulting in large sac-like alveoli^58^. Such inhibition of alveolar septation suggests an important role for thyroid hormone in the postnatal structural maturation of the lungs^58^. It was also reported that during early embryonic lung development, THs accelerate the epithelial and mesenchymal cell differentiation at the expense of branching morphogenesis and lung growth^58,62^. We found that the transcript levels of the *angptl3* that participates in the alveolarization during postnatal lung development in mammals, are increased at Stage IV of metamorphosis. This suggests that alveolarization may occur in the lungs during the final stage of metamorphosis, similarly to what occurs before and soon after birth in mammals.

During metamorphosis, the aquatic larvae of *A. tigrinum* carry out the gas exchange of O_2_ and CO_2_ mainly by the cutaneous and branchial way and, to a smaller proportion, by the lungs^64^. However, by the end of metamorphosis the branchial tissue has disappeared and terrestrial organisms obtain up to 60% of O_2_ through the metamorphic pulmonary system^64^. Branchial tissue progressive degradation during the metamorphosis process of *A. mexicanum* has been related to the increase in expression of genes whose function has been associated with apoptosis, such phenomenon correlates with the increase in expression of genes involved in developmental changes in the epidermis^24^. According to our results in the enriched cluster genes for gills we identified *Avpkp1* whose ortholog has been involved in the formation of stratified epithelium during embryonic development in mammals^49^, while in the cluster of gradually decreasing genees in this tissue we identified *rhcg*, a gene that, in aquatic organisms, modulates the excretion of ammonium^51^. This could be related to the change of aquatic to amphibian habits, due to the fact that aquatic organisms excrete a greater amount of ammonium and less urea. As a result of the changes of habitats the skin also undergoes modifications to adapt to the new environment as previously described for *A. mexicanum.* In correlation with such changes, we found that *Avhoxa7* transcripts increase in the gills transcriptome. HOXA7 expresses in suprabasal fetal human epidermis, with expression persisting in the adult epidermis but not in the dermis, correlating with its function in modulating the expression of keratin genes such as *Avkrt7*^52^. The process of keratinization of the skin allows organisms to modulate environmental dehydration retaining moisture^51^.

In conclusion, the profiling of the transcriptional programs and the respective molecular pathways underlying the morphophysiological changes in the circulatory and respiratory systems during metamorphosis, can help to better understand the complex developmental processes that allow amphibians to transit from aquatic to a terrestrial environment. Some of the pathways described as changing in heart and lung during *A. velasci* metamorphosis are related to pre and postnatal morphological events in mammals, both under normal developmental conditions as well as in human diseases associated with aberrant development and function of lung and heart. Finally, our work represents the first transcriptional landscape along the diverse stages of the metamorphosis of a viable metamorphic *Ambystoma spp*. This study becomes the foundation for the immediate generation of biological questions and their experimental test at organ, tissue, cellular and molecular levels. Such future experimentally approaches will shed light on the morphophysiological and molecular events that govern not only the metamorphosis phenomenon, but also heart and lung proper development and maturation.

## Acknowledgements

The authors acknowledge Biol. Arturo Vergara-Iglesias from (CIBAC-UAM Xochimilco) and Ollin Olivia Ramírez from PYMVS Ambystomania for support in animal sampling and management. We also thank UMA San Miguel Tecocomulco for reproduction and conservation efforts. ACR wishes to dedicate this manuscript to Tepeapulco, Hidalgo; México, the home of *Ambystoma velasci.*

## Author Contributions

Conceived the study: JPM and ACR. Designed the experiments: JPM, ACR, JCP, AEC, ESV, HL and SD. Performed the experiments: JPM, AEC, ESV, MCG and SD. Analyzed the data: JPM, JCP, ACR, AEC, JCM and MCG. Contributed reagents/materials/analysis tools: ACR, ESV, HL, JCM and SD. Wrote the paper: JPM and ACR with inputs from all co-authors.

## Data availability

The datasets generated during the current study will be available under an NCBI BioProject,which will be shared once tha manuscript is accepted for publication in a regular journal.

Supplementary tables, video, (cited along the text) and other data will be available once this manuscript is accepted.

## References

1. Laudet, V. The origins and evolution of vertebrate metamorphosis. Current Biology. 21, 726–737, doi:10.1016/j.cub.2011.07.030 (2011).

2. Wang, X. et al. Thyroid hormone-responsive genes mediate otolith growth and development during flatfish metamorphosis. Comparative Biochemistry and Physiology Part A. 158, 163–168. doi:10.1016/j.cbpa.2010.10.014 (2011).

3. Ferraresso, S. et al. Exploring the larval transcriptome of the common sole (*Solea solea* L.). BMC Genomics. 14, 315, doi:10.1186/1471-2164-14-315 (2013).

4. Zhao, L., Liu, L., Wang, S., Wang, H., & Jiang, J. Transcriptome profiles of metamorphosis in the ornamented pygmy frog Microhyla fissipes clarify the functions of thyroid hormone receptors in metamorphosis. Scientific reports. 6, 27310, doi:10.1038/srep27310 (2016).

5. Liu, L., Zhu, W., Liu, J., Wang, S., & Jiang, J. Identification and differential regulation of microRNAs during thyroid hormone-dependent metamorphosis in *Microhyla fissipes*. BMC genomics. 19, 507, doi:10.1186/s12864-018-4848-x (2018).

6. Tata, J. Hormonal signalling during amphibian metamorphosis. Proceedings of the Indian National Sciences Academy. 69, 773–790 (2003).

7. Holstein, T., & Laudet, V. Life-History Evolution: At the Origins of Metamorphosis. Current Biology. 24, 159–161, doi:10.1016/j.cub.2014.01.003 (2014).

8. Brown, D. D., & Cai, L. Amphibian metamorphosis. Developmental biology. 306, 20–33, doi:10.1016/j.ydbio.2007.03.021 (2007).

9. Furlow, J., Berry, D., Wan, Z., & Brown, D. A set of novel tadpole specific genes expressed only in the epidermis are down-regulated by thyroid hormone during *Xenopus laevis* metamorphosis. Development biology. 182, 284–298, doi:10.1006/dbio.1996.8478 (1997).

10. Ishizuya-Oka, A., Hasebe, T., & Shi, Y. B. Apoptosis in amphibian organs during metamorphosis. Apoptosis: an international journal on programmed cell death. 15, 350–64, doi:10.1007/s10495-009-0422-y (2010).

11. Das, B. et al. Gene expression changes at metamorphosis induced by thyroid hormone in *Xenopus laevis* tadpoles. Developmental Biology. 291, 342–355 doi:10.1016/j.ydbio.2005.12.032 (2006).

12. Ohmura, H., & Wakahara, M. Transformation of skin from larval to adult types in normally metamorphosing and metamorphosis-arrested salamander, Hynobius retardatus. Differentiation. 63, 237–246, doi:10.1046/j.1432-0436.1998.6350237.x (1998).

13. Hasebe, T., Kajita, M., Shi, Y. B., & Ishizuya-Oka, A. Thyroid hormone-up-regulated hedgehog interacting protein is involved in larval-to-adult intestinal remodeling by regulating sonic hedgehog signaling pathway in Xenopus laevis. Developmental dynamics: an official publication of the American Association of Anatomists. 237, 3006–15, doi:10.1002/dvdy.21698 (2008).

14. Damjanovski, S., Puzianowska-Kuznicka, M., Ishizuya-Oka, A., & Shi, Y. B. Differential regulation of three thyroid hormone-responsive matrix metalloproteinase genes implicates distinct functions during frog embryogenesis. FASEB J. 14, 503–510, doi:10.1096/fasebj.14.3.503 (2000).

15. Shi, Y., Fu, L., Hsia, S., Tomita, A., & Buchholz, D. Thyroid hormone regulation of apoptotic tissue remodeling during anuran metamorphosis. Cell Research. 11, 245–252, doi:10.1038/sj.cr.7290093 (2001).

16. Mukhi, S., Cai, L., & Brown, D. Gene switching at *Xenopus laevis* metamorphosis. Developmental biology. 338, 117–126, doi:10.1016/j.ydbio.2009.10.041 (2010).

17. Johnson, C., & Voss, R.S. Salamander paedomorphosis:linking thyroid hormone to life history and life cycle evolution. Current Topics in Developmental Biology. 103, 229–258, doi:10.1016/B978-0-12-385979-2.00008-3 (2013).

18. Buchholz, D., & Hayes, T. Variation in thyroid hormone action and tissue content underlies species differences in the timing of metamorphosis in desert frogs. Evolution and Development. 7, 458–467, doi:10.1111/j.1525-142X.2005.05049.x (2005).

19. Larras-Regard, E., Taurog, A., & Dorris, M. Plasma T4 and T3 levels in *Ambystoma tigrinum* at various stages of metamorphosis. General and Comparative Endocrinology. 4, 443–450, doi:10.1016/0016-6480(81)90228-8 (1981).

20. Cano-Martínez, A., Vargas-Gonzalez, A., & Asai, M. Metamorphic stages in *Ambystoma mexicanum*. Axolotl Newsletter. 23, 64–71 (1994).

21. Demircan, T. et al. A histological atlas of the tissues and organs of neotenic and metamorphosed axolotl. Acta Histochemica. 118, 746–759, doi: 10/1016/j.achthis.2016.07.006 (2016).

22. Badawy, G. Effect of thyroid stimulating hormone on the ultrastructure of the thyroid gland in the Mexican axolotl during metamorphic climax. Journal of Applied Pharmaceutical Science. 1, 60–66 (2011).

23. Voss, S., Kump, D., Walker, J., Shaffer, H., & Voss, G. Thyroid hormone responsive QTL and evolution of paedomorphic salamanders. Heredity. 109, 293–298, doi:10.1038/hdy.2012.41 (2012).

24. Page, R. et al. Effect of thyroid hormone concentration on the transcriptional response underlying induced metamorphosis in the mexican axolotl (*Ambystoma mexicanum*). BMC Genomics. 9, 78, doi:10.1186/1471-2164-9-78 (2008).

25. Page, R. B., Monaghan, J. R., Walker, J. A., & Voss, S. R. A model of transcriptional and morphological changes during thyroid hormone-induced metamorphosis of the axolotl. General and comparative endocrinology. 162, 219–32, doi:10.1016/j.ygcen.2009.03.001 (2009).

26. Huggins, P. et al. Identification of differentially expressed thyroid hormone responsive genes from the brain of the Mexican Axolotl (*Ambystoma mexicanum*). Comparative biochemistry and physiology. Toxicology & pharmacology: CBP. 155, 128–35, doi:10.1016/j.cbpc.2011.03.006 (2012).

27. Garg, V. et al. GATA4 mutations cause human congenital heart defects and reveal an interaction with TBX5. Nature. 424, 443–447, doi:10.1038/nature01827 (2003).

28. Cui, Y. et al. Single-cell transcriptome analysis maps the developmental track of the human heart. Cell Reports. 26, 1934–1950, doi:10.1016/j.celrep.2019.01.079 (2019).

29. Wang, J. et al. Targeted disruption of *Smad4* in cardiomyocytes results in cardiac hypertrophy and heart failure. Circulation research. 97, 821–828, doi:10.1161/01.RES.0000185833.42544.06 (2005).

30. McDouall, R. M., Puspa, B., McCormack, A., Yacoub, M. H., & Rose, M. L. MHC Class II expression on human heart microvascular endothelial cells: Exquisite sensitivity to interferón-gand natural killer cells. Transplantation. 64, 1175–1180, doi:10.1097/00007890-199710270-00016 (1997).

31. Chistiakov, D. A., Melnichenko, A. A., Myasoedova, V. A., Grechko, A. V., & Orekhov, A. N. Thrombospondins: A role in cardiovascular disease. International Journal of Molecular Sciences. 18, 1540, doi:10.3390/ijms18071540 (2017).

32. Akazawa, H., & Komuro, I. Cardiac transcription factor Csx/Nkx2-5: Its role in cardiac development and diseases. Pharmacology & Therapeutics, 107, 252–268, doi:10.1016/j.pharmthera.2005.03.005 (2005).

33. Cantú, I. & Philipsen, S. Flicking the switch: adult hemoglobin expression in erythroid cells derived from cord blood and human induced pluripotent stem cells. Haematologica. 99, 1647–1649, doi:10.3324/haematol.2014.116483 (2014).

34. Sahara, M., Santoro, F., & Chien, K. R. Programming and reprogramming a human heart cell. The EMBO Journal. 34, 710–738, doi: 10.15252/embj.201490563 (2015).

35. England, J., & Loughna, S. Heavy and light roles: myosin in the morphogenesis of the heart. Cellular and molecular life sciences: CMLS. 70, 1221–39, doi:10.1007/s00018-012-1131-1 (2013).

36. Geng, L. et al. Syntaxin 5 regulates the endoplasmic reticulum channel-release properties of polycystin-2. PNAS. 105, 15920–15925, doi:10.1073/pnas.0805062105 (2008).

37. Yang, J. N., Wang, Y., Garcia-Roves, P. M., Björnholm, M., & Fredholm, B. Adenosine A(3) receptors regulate heart rate, motor activity and body temperature. Acta physiologica (Oxford, England). 199, 221–30, doi: 10.1111/j.1748-1716.2010.0209.x (2010).

38. Anumonwo, J. M., & Lopatin, A. N. Cardiac strong inward rectifier potassium channels. Journal of molecular and cellular cardiology. 48, 45–54, doi:10.1016/j.yjmcc.2009.08.013 (2010).

39. Wei, L. et al. Inhibition of Rho family GTPases by Rho GDP dissociation inhibitor disrupts cardiac morphogenesis and inhibits cardiomyocyte proliferation. Development. 129, 1705–1714 (2002).

40. Yang, Y. T., Rayburn, H., & Hynes, R. Cell adhesión events mediates by a4 integrins are essential in placental and cardiac development. Development. 121, 549–560 (1995).

41. Arora, R., Metzger, R. J., & Papaioannou. Multiple roles and interactions of *Tbx4* and *Tbx5* in development of the respiratory system. PLOS Genetics. 8, e1002866, doi:10.1371/journal.pgen.1002866 (2012).

42. Liu, H. et al. HOXC8 promotes proliferation and migration through transcriptional up-regulation of TGFb1 in non-small cell lung cancer. Oncogenesis. 7, 1–15, doi:10.1038/s41389-017-0016-4 (2018).

43. Choi, K. et al. Distinct biological roles for the Notch ligands Jagged-1 and Jagged-2. The journal of biological chemistry. 284, 17766–17774, doi:10.1074/jbc.M109.003111 (2009).

44. Portnoy, A. et al. Effects of urotensin II receptor antagonist, GSK1440115, in asthma. Frontiers in pharmacology. 4, 1–7, doi:10.3389/fphar.2013.00054 (2013).

45. Zhang, Y. et al. PR-domain-containing *Mds-Evil*in critical for long-term hematopoietic stem cell function. Blood. 118, 3853–3861, doi:10.1182/blood-2011-02-334680 (2011).

46. Lai, C. K. et al. Functional characterization of putative cilia genes by high-content analysis. Molecular biology of the cell. 22, 1105–1119, doi:10.1091/mbc.E10-07-0596 (2011).

47. Ma, Y. J, Lee, B. L., & Garred, P. An overview of the synergy and crosstalk between pentraxins and collectins/ficolins: their functional relevance in complement activation. Experimental & Molecular Medicine. 49, e320, doi:10.1038/emm.2017.51 (2017).

48. Mammoto, T. et al. LRP5 regulates development of lung microvessels and alveoli through the angiopoietin-Tie2 pathway. PLoS One. 7, e41596, doi:10.1371/journal.pone.0041596 (2012).

49. McGrath, J. A. et al. Mutations in the plakophilin 1 gene result in ectodermal dysplasia/skin fragility syndrome. Nature genetics. 17, 240–244, doi:10.1038/ng1097-240 (1997).

50. He, J., & Baum, L. G. Presentation of Galectin-1 by Extracellular Matrix triggers T cell death. The Journal of Biological Chemistry. 279, 4705–4712, doi:10.1074/jbc.M311183200 (2004).

51. Evans, D. H., Piermarini, P. M., & Choe, K. P. The multifunctional fish gill: dominant site of gas exchange, osmoregulation, acid-base regulation, and excretion of nitrogenous waste. 120–138, doi:10.1152/physrev.00050.2003 (Physiology Reviews, 2005).

52. La Celle, P. T., & Polakowska, R. R. Human homeobox HOXA7 regulates keratinocyte Transglutaminase Type 1 and inhibits differentiation. The Journal of Biological Chemistry. 276, 32844–32853, doi:10.1074/jbc.M104598200 (2001).

53. Sandilands, A. et al. Generation and characterisation of Keratin 7 (K7) knockout mice. PLoS ONE. 8, e64404, doi:10.1371/journal.pone.0064404 (2013).

54. Kelwick, R., Desanlis, I., Wheeler, G. N., & Edwards, D. R. The ADAMTS (A Disintegrin and Metalloproteinase with Thrombospondin motifs) family. Genome Biology. 16:113, doi: 10.1186/s13059-015-0676-3 (2015).

55. Brent, G. A. Mechanisms of thyroid hormone action. The Journal of clinical investigation. 122, 3035–43, doi:10.1172/JCI60047 (2012).

56. Granados-Riveron, J. T. et al. α-Cardiac myosin heavy chain (*MYH6*) mutations affecting myofibril formation are associated with congenital heart defects. Human Molecular Genetics. 19, 4007–4016, doi:10/10.93/hmg/ddq315 (2010).

57. Atlas, S. A. The Renin-Angiotensin Aldosterone System: pathophysiological role and pharmacologic inhibition. Supplement to Journal of Managed Care Pharmacy. 13, 9–20, doi:10.18553/jmcp.2007.13.s8-b.9 (2007).

58. Tuyl, M. et al. Prenatal exposure to thyroid hormone is necessary for normal postnatal development of murine heart and lungs. Developmental Biology. 272, 104–117, doi:10.1016/j.ydbio.2004.03.042 (2004).

59. Hoffman, A. D., Peterson, M. A., Friedland-Little, J. M., Anderson, S. A. & Moskowitz, I. P. Sonic hedgehog is required in pulmonary endoderm for atrial septation. Development. 136, 1761–1770, doi:10.1242/dev.034157 (2009).

60. Mohammadi, M. M. et al. The transcription factor GATA4 promotes myocardial regeneration in neonatal mice. EMBO Molecular Medicine. 9, 265–279, doi:10.15252/emmm.201606602 (2017).

61. Xu, H. et al. Tbx1 has a dual role in the morphogenesis of the cardiac outflow tract. Development. 131, 3217–3227, doi:10.1242/dev.01174 (2004).

62. Chung, C. et al. Hippo-Foxa2 signaling pathway plays a role in peripheral lung maturation and surfactant homeostasis. PNAS. 110, 7732–7737, doi:10.1073/pnas.1220603110 (2013).

63. Li, M. et al. Thyroid hormone action in postnatal heart development. Stem Cell Research. 13, 582–591, doi:10.1016/j.scr.2014.07.001 (2014).

64. Burggren, W. W. The respiratory transition from water to air breathing during amphibian metamorphosis. American Zoologist, 34, 238–246 (1994).

65. Zachary, R. L., & Hanken, J. Convergent evolutionary reduction of atrial septation in lungless salamanders. Journal of Anatomy. 230, 16–29, doi:10.1111/joa.12535 (2017).

66. Coots, P. S., & Seifert, A. W. 2015. Thyroxine-induced metamorphosis in the axolotl (*Ambystoma mexicanum*). Salamanders in Regeneration Research: Methods and Protocols, Methods in Molecular Biology. Springer. Vol 1290. doi: 10.1007/978-1-4939-2495-0_11.

67. Grabherr, M. G. et al. Trinity: reconstructing a full-length transcriptome without a genome from RNA-Seq data. Nature Biotechnology, 29, 644–652, doi:10.1038/nbt.1883 (2013).

68. Bray, N. L., Pimentel, H., Melsted, P., & Pachter, L. Near-optimal probabilistic RNA-seq quantification. Nature Biotechnology. 34, 525–527, doi:10.1038/nbt.3519 (2016).

69. Mi, H. et al. Protocol Update for large-scale genome and gene function analysis with the PANTHER classification system (v.14.0). Nature Protocols. 8, 1551–1566, doi:10.1038/nprot.2013.092 (2019).

