## Supplemental Figures for "Multi-organ transcriptomic landscape of *Ambystoma velasci* metamorphosis"

[1]*Molecular & Developmental Complexity Group,* Unit of Advanced Genomics, UGA-CINVESTAV, Irapuato, México.

[2]Department of Biochemistry, Escuela Nacional de Ciencias Biológicas, Instituto Politécnico Nacional, México City, México.

[3]Departamento de Genética del Desarrollo y Fisiología Molecular, Instituto de Biotecnología, Universidad Nacional Autónoma de México, AP 510-3, Cuernavaca, Mor. 62250, México.

[4]Departamento de Medicina Genómica y Toxicología Ambiental, Instituto de Investigaciones Biomédicas, Universidad Nacional Autónoma de México, Ciudad Universitaria, México DF 04510, México.

### Supplementary

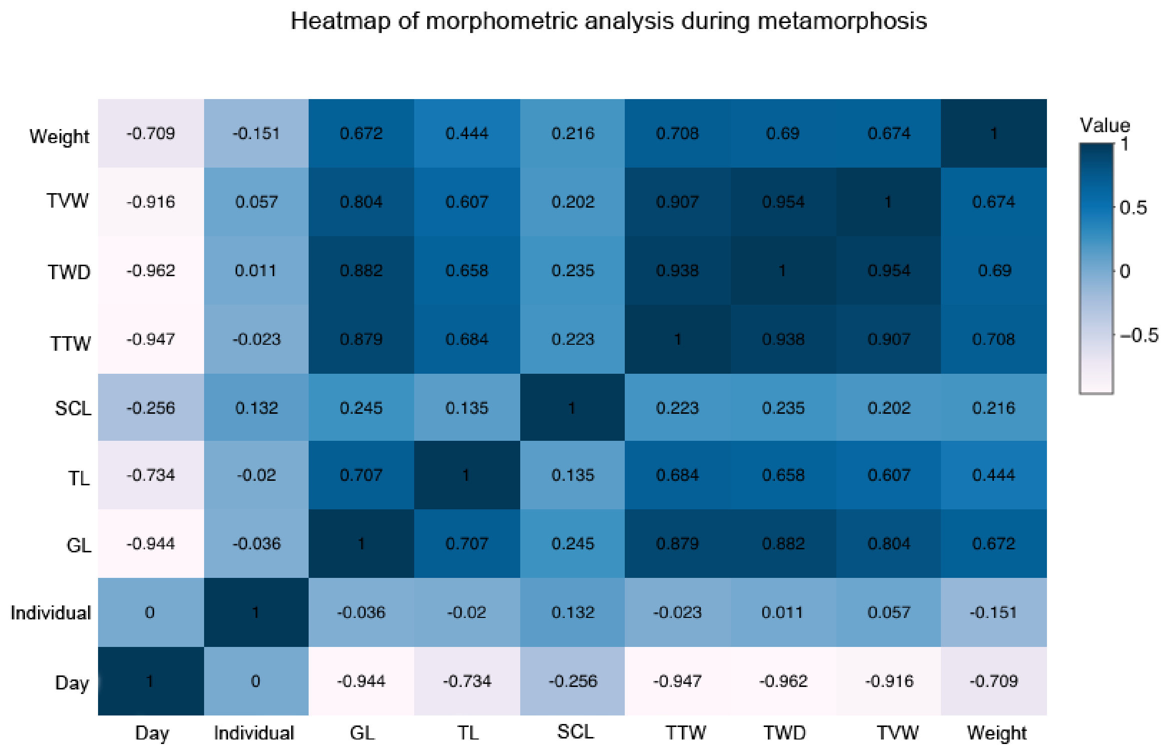


**SFigure 1. Multiple Correlation Analysis.** This analysis represents the correlation between the variables total length (TL), snout-cloaca length (SCL), total tail width (TTW), tail dorsal width (TDW), tail ventral

width (TVW), gill length (GL) and weight (g) in metamorphosis process. The variables GL, TDW, and TTW have a greater correlation with metamorphosis ($p<0.05$).

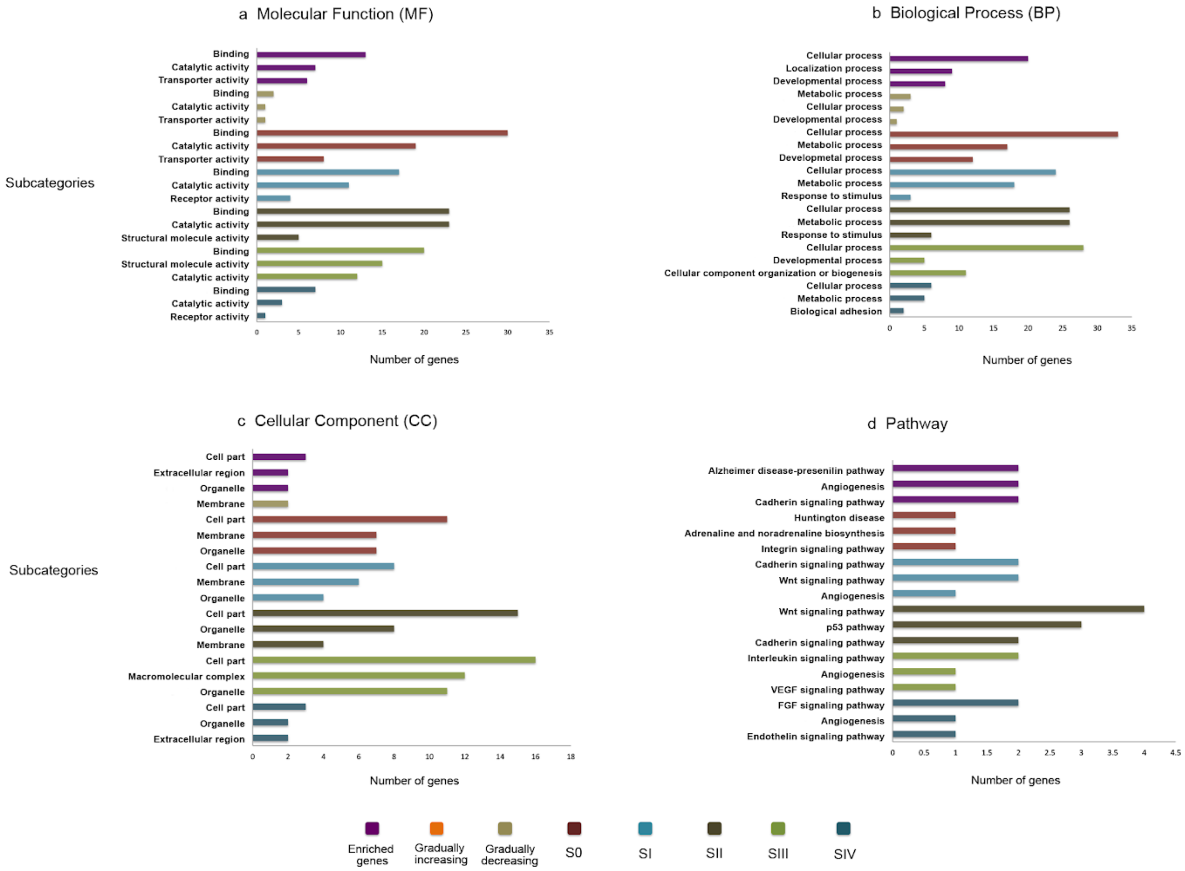


**SFigure 2. Gene Ontology analysis in lung during metamorphosis.** Enriched functional categories of DEGs in lung by expression pattern during metamorphosis. Stage 0 is used as reference. In the Y axis the subcategories of the gene ontology are represented, in the X axis the number of genes enriched in each subcategory. The (**a**) represents Molecular Function category (MF), (**b**) Biological Process (BP), (**c**) Cellular Component (CC) and (**d**) Pathway. The clusters are represented by different colors: purple for heart-enriched genes compared to lung and gills, orange for gradually increased expression, red for genes enriched in S0, blue for SI-12 hpi, brown for SII-36 hpi, green for SIII 6-12 dpi and turquoise for SIV 17-23 dpi.

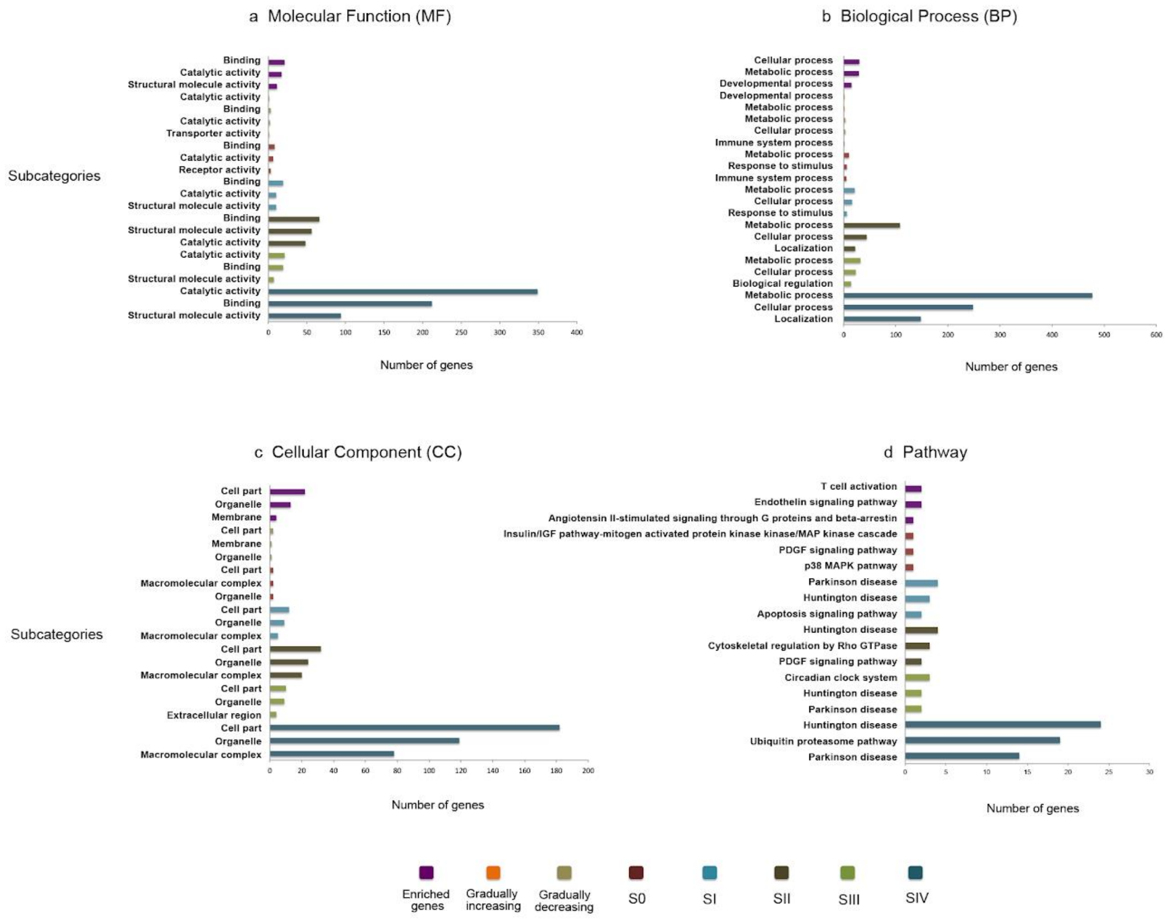


**SFigure 3. Gene Ontology analysis in gills during metamorphosis.** Enriched functional categories of DEGs in gills by expression pattern during metamorphosis. Stage 0 is used as reference. In the Y axis the subcategories of the gene ontology are represented, in the X axis the number of genes enriched in each subcategory. The (**a**) represents Molecular Function category (MF), (**b**) Biological Process (BP), (**c**) Cellular Component (CC) and (**d**) Pathway. The clusters are represented by different colors: purple for heart-enriched genes compared to lung and gills, orange for gradually increased expression, red for genes enriched in S0, blue for SI-12 hpi, brown for SII-36 hpi, green for SIII 6-12 dpi and turquoise for SIV 17-23 dpi.

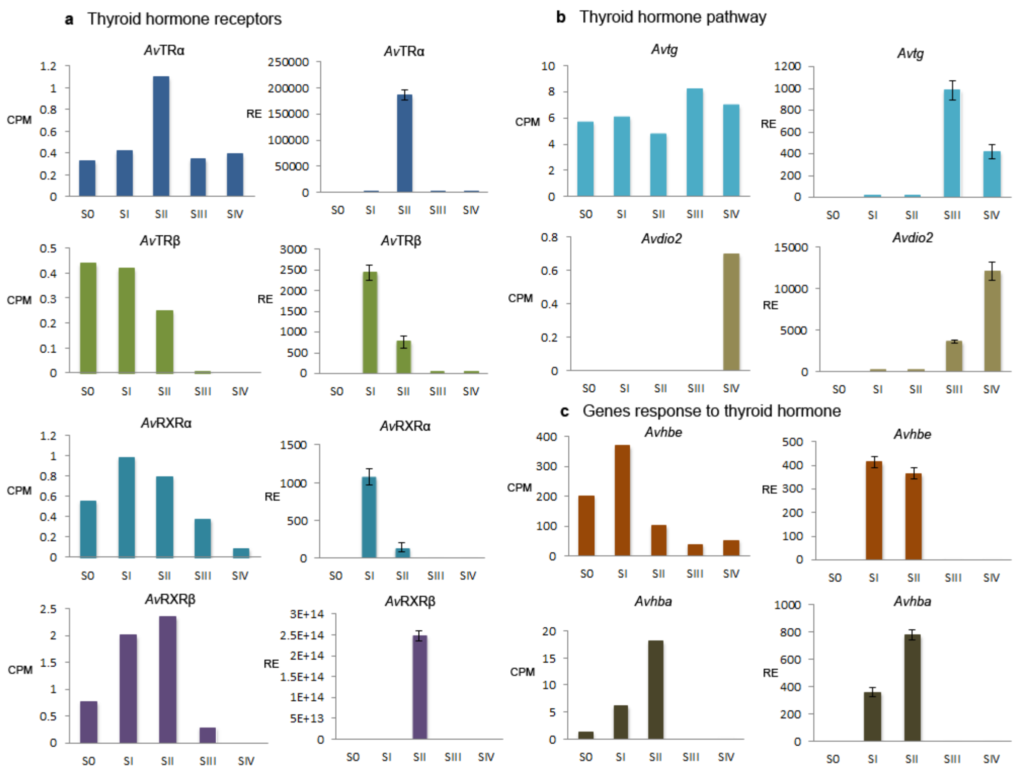


**SFigure 4. qRT-PCR validation for TH pathway in lung.** The X axis represents the metamorphosis stages S0, SI-12 hpi, SII-36 hpi, SIII 6-12 dpi and SIV 17-23 dpi. The counts per million (CPM) value represents the transcriptome result. The relative expression (RE) is the qPCR analysis, the columns and bars represent the means and the standard deviation of the three individual samples. (**a**) Expression patterns of *A. velasci* homologous genes for thyroid hormone receptors: *Av*TRα, *Av*TRβ, *Av*RXRα and *Av*RXRβ (**b**) Genes involved in thyroid hormone synthesis, *Avtg* and *Avdio2*. (**c**) Genes that have been reported as responsive to thyroid hormone; *Avhbe* and *Avhba*.

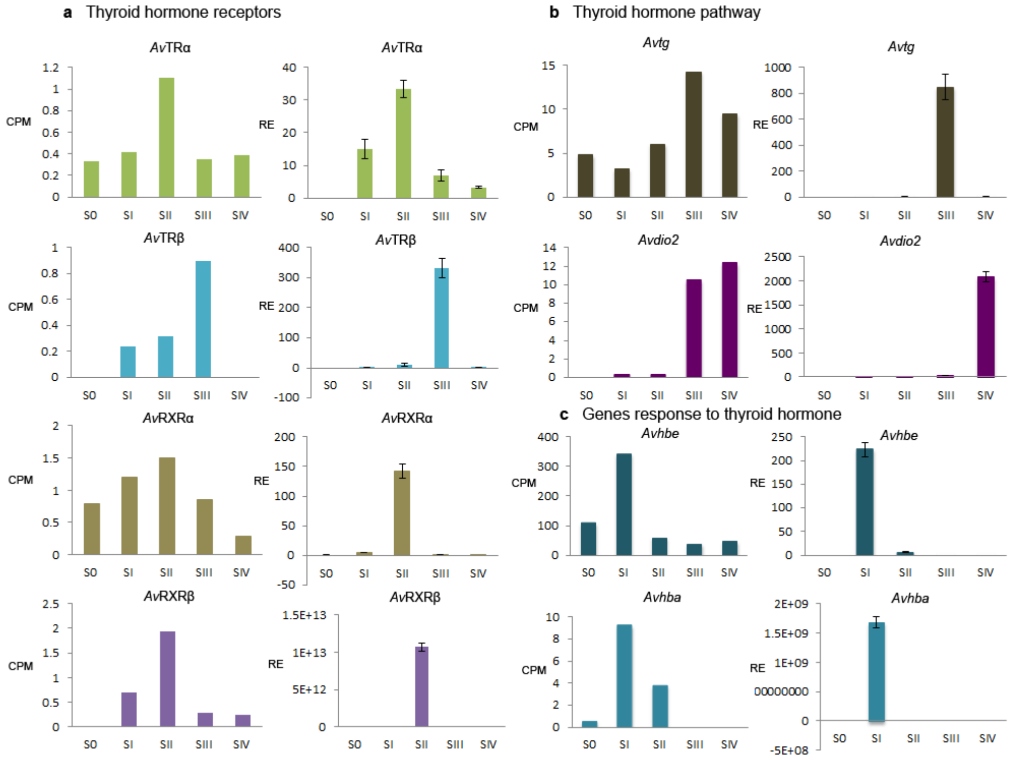


**SFigure 5. qRT-PCR validation for TH pathway in gills.** The X axis represents the metamorphosis stages S0, SI-12 hpi, SII-36 hpi, SIII 6-12 dpi and SIV 17-23 dpi. The counts per million (CPM) value represents the transcriptome result. The relative expression (RE) is the qPCR analysis, the columns and bars represent the means and the standard deviation of the three individual samples. (**a**) Expression patterns of *A. velasci* homologous genes for thyroid hormone receptors: *Av*TRα, *Av*TRβ, *Av*RXRα and *Av*RXRβ (**b**) Genes involved in thyroid hormone synthesis, *Avtg* and *Avdio2*. (**c**) Genes that have been reported as responsive to thyroid hormone; *Avhbe* and *Avhba*.
